# Unsupervised Machine Learning for Data Encoding applied to Ovarian Cancer Transcriptomes

**DOI:** 10.1101/855593

**Authors:** Tom M George, Pietro Lio

## Abstract

Machine learning algorithms are revolutionising how information can be extracted from complex and high-dimensional data sets via intelligent compression. For example, unsupervised Autoen-coders train a deep neural network with a low-dimensional “bottlenecked” central layer to reconstruct input vectors. Variational Autoencoders (VAEs) have shown promise at learning meaningful latent spaces for text, image and more recently, gene-expression data. In the latter case they have been shown capable of capturing biologically relevant features such as a patients sex or tumour type. Here we train a VAE on ovarian cancer transcriptomes from The Cancer Genome Atlas and show that, in many cases, the latent spaces learns an encoding predictive of cisplatin chemotherapy resistance. We analyse the effectiveness of such an architecture to a wide range of hyperparameters as well as use a state-of-the-art clustering algorithm, t-SNE, to embed the data in a two-dimensional manifold and visualise the predictive power of the trained latent spaces. By correlating genes to resistance-predictive encodings we are able to extract biological processes likely responsible for platinum resistance. Finally we demonstrate that variational autoencoders can reliably encode gene expression data contaminated with significant amounts of Gaussian and dropout noise, a necessary feature if this technique is to be applicable to other data sets, including those in non-medical fields.

## 1 Introduction

The development of gene expression profiling, a technique allowing scientists to measure the expression levels of thousands of genes at once [1, 2, 3, 4], has led to biomedical research becoming ever more data-intensive. Large-scale genomic efforts (including, but not limited to, cancer research) are mapping the genetic and biochemical alterations which cause cancer, yet interpreting and translating these data into treatments remains hard[5]. This is compounded by the fact that cancer is a dynamic disease which can move unpredictably to other tissues through a process known as metastasis. In the words of Robert Austin (Princeton Professor of Biophysics) “cancer needs more big ideas - and those of scientists from other disciplines should be taken more seriously”[6]. Specifically disciplines such as physics, data science and mathematics. From using soap bubbles to model cell division[7] to studying the mathematics of genes mutation rates[8, 9], there is a rich history of the physical sciences contributing to cancer research and most experts agree that the continued fusion of these disciplines with cancer biology is the most likely route towards a deeper understanding of, and potentially a cure for, cancer in future years[10]. In this report we propose that unsupervised machine learning could be used to shed light on poorly understood processes relating to ovarian cancer.

The study of machine learning has flourished over the past decade largely due to an exponential increases in computing power. Machine learning is renowned for its ability to learn complex input-output relationships in fields where data sets are large and complex - from theoretical physics to biomedicine. For example, in high energy physics machine learning has the potential to assist with event classification at the LHC [11, 12], whilst in astrophysics neural networks are already routinely used for galaxy classification[13]. In more applied physical sciences, neural networks could assist climate scientists by predicting oceanic heat flows from satellite imagery [14, 15]. In general, machine learning proves most fruitful when the ability of a field to collect data has outpaced its ability to analyse it. Nowhere is this more pertinent than in the field of biomedicine.

In this project we study how variational autoencoders (a relatively new class of machine learning algorithms, see section 2.2) can be applied to a data set of ovarian cancer transcriptomes in order to learn about the biological mechanisms behind *platinum chemotherapy resistance*. Platinum (or cisplatin) resistance is a phenomenon where ovarian cancer patients fail to respond to platinum based chemotherapy (currently first-line therapy for ovarian cancer[16]) - they transition from being ‘platinum sensitive’ to ‘platinum resistant’. The high mortality rate[17, 18, 19] associated with ovarian cancer is, in part, due to our lack of understanding about this aspect. Here we use a variational autoencoder (VAE) to compress high-dimensional ovarian cancer transcriptomes into a biologically meaningful latent space and show it is possible for one or more of the encodings to learn a useful representation of platinum resistance. Whilst others have shown VAEs can learn to represent pronounced biological feature, like tumour type[20] or patient sex[21], to the best of our knowledge we are the first to extend this to much harder task of encoding platinum resistance.

We comprehensively analyse how the VAE hyperparameters affect encoding performance as a tool for anyone looking to applying this methodology further. Enrichment analysis is performed to retrace our steps and convert the meaningful information stored in the latent variables into a useful list of biological process associated with platinum resistance. Finally, we investigate how robust the methodology is to data artificially contaminated with noise.

Lying firmly at the interface between data science and cancer research, this report would benefit from a comprehensive literature review covering the history and state-of-the-art methods in both fields. Unfortunately due to the lack of space this will not be possible however section 2 pays particular attention to the theory and mathematics behind the machine learning we use here. I believe the theoretical concepts discussed in this section to be the most far reaching^1^ and interesting to an audience of physicists - after all this work was undertaken from a physics background.

## 2 Theory

Machine learning is a branch of computer science where models are iteratively trained to learn a mapping between two types of data. Artificial neural networks (ANNs) takes the form of multiple connected layers of ‘neurons’ whose values are derived through a non-linear map from the previous layer via a trainable weight matrix, figure 1. Neural networks are powerful tools, outperforming humans at many tasks such as supervised image classification and pattern recognition[22, 23]. A basic understanding of deep learning (specifically ANNs, gradient descent, loss functions and activation functions) will be assumed from hereon however for some introductory reading I suggest this eBook by Michael Neilson: ‘Neural Networks and Deep Learning’ [24].

**Figure 1:**
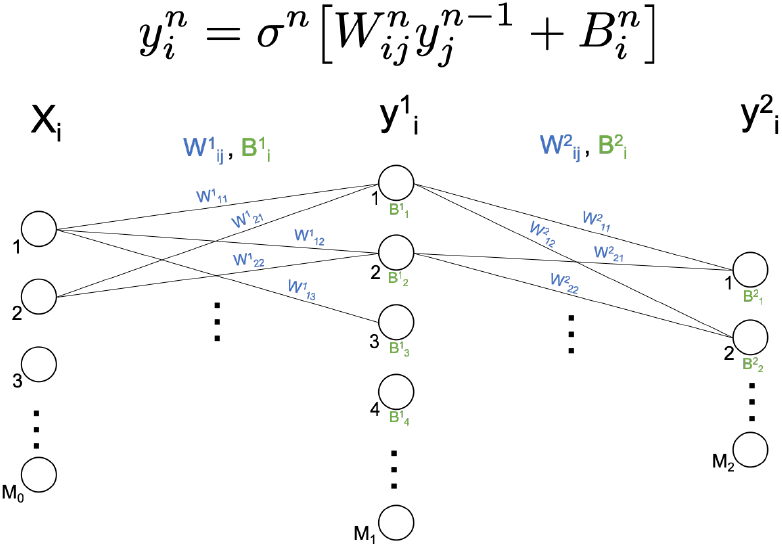
A two layer fully connected artificial neural network (ANN). Information feeds forward from the input vector *X* to the output vector *y* according to the equation shown. The weights and biases are successively trained by backpropagation to reduce a chosen ‘loss function’.

### 2.1 Autoencoders

An autoencoder is a neural network trained to reconstruct the original input rather than a classification label (**x** → **x** rather than **x** → **y**). One benefit of unsupervised over supervised machine learning is that it can be used to learn about large and complex data sets which have not necessarily been painstakingly labelled. The reconstruction task is made difficult (and thus non-trivial) by the constraint that an internal hidden layer, **z**, is designed to have a dimension far smaller than the input. This “bottleneck” forces the autoencoder to compress the data (**z** is also known as the ‘code’ or the ‘latent vector’) by retaining only the most essential information needed to reproduce the input, see figure 2.

**Figure 2:**
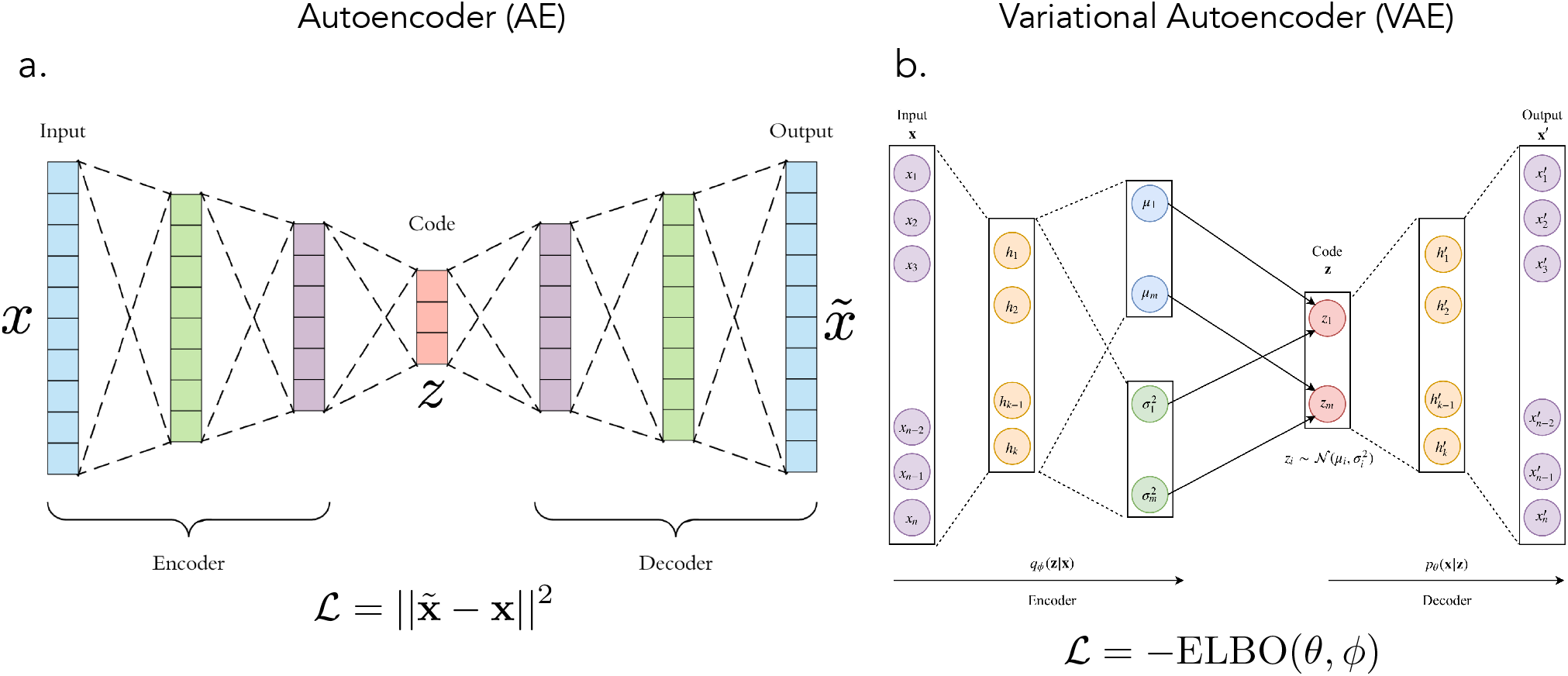
(a) Schematic of an autoencoder (AE). The data is compressed into a low dimension latent space **z** before being reconstructed again. The loss function is usually the euclidean reconstruction error. Figure source [27]. (b) Schematic of a variational autoencoder (VAE). The VAE learns the parameters of a conditional probability distribution *q_ϕ_* which is then sampled and decoded. The loss function (negative ELBO, see equation (3)) encourages the VAE to learn to reconstruct the data after compression but also to learn a latent representation of the data which is smooth and Gaussian. Figure source [28].

The *encoder* maps the input vector to it’s latent representation: *enc*: 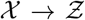. The *decoder* transforms 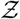 back to the input vector space, *dec*: 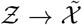. The autoencoder is trained using a gradient descent algorithm to minimise the reconstruction loss:

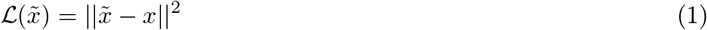

where 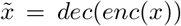. The most obvious use for autoencoders is for dimensionality reduction (e.g. data compression) however they have also been applied successfully to tasks from natural language processing [25] to the prediction of new high-energy physics [26].

### 2.2 Variational Autoencoders

An exciting variant of the autoencoder is the *variational autoencoder*, initially proposed by Kingma and Welling [29]. VAEs are trained with the additional constraint that they must generate latent vectors with a Gaussian distribution. Instead of learning deterministic mapping functions, the encoder and decoder now learn parameters of probability distribution functions that model the training data [30, 31].

Suppose we have some data set which in its native (high-dimensional) vector space can be represented by a probability distribution function *p*(**x**), a function *predetermined* by the data set. We wish to represent this data set in a low dimensional space with a latent variable which has a prior distribution *p*(**z**), a function we *choose*. The process for generating data is to sample a latent vector from *p*(**z**) and then form the parameterised conditional probability distribution *p_θ_*(**x**|**z**) from which we can sample **x**. The conditional probability function is ‘good’ if it maximises

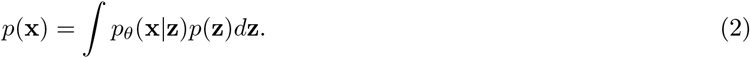

evaluated for each **x** vector in the training data set i.e. *p_θ_*(**x**|**z**) is likely to generate data points close to those in our data set. Unfortunately, calculating this is hard since it involves integrating over the whole space of **z** values which is often large. A Monte Carlo attempt, 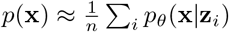 for **z** sampled from *p*(**z**), will fail since even if we initially pick *p_θ_*(**x**|**z**) to be a good generator, the input dimension is so high that the vast majority of samples are unlikely to produced anything like **x** so the integral will <u>not</u> converge[32].

To surpass this we cheat and, instead of sampling **z** from *p*(**z**) we sample it from *q_ϕ_*(**z**|**x**). *q_ϕ_* is trained to assign high probability values to **z**’s that are likely to have created **x** - exactly what we need for convergence. Now, instead of maximising 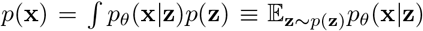, we’ll be maximising 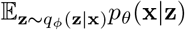. The relationship between these two quantities is a cornerstone of variational bayesian methods [33]. The difference is encapsulated in the following equation for the ‘Evidence Lower Bound Objective (ELBO)’, derived in appendix A,

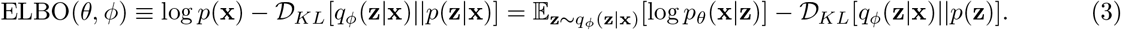

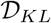 is the Kullback-Leibler divergence and quantifies the difference between two functions, vanishing if they are identical. The ELBO therefroe contains exactly what we want to maximise, i.e. *p*(**x**) along with an error term which will become small if *q_ϕ_* is high capacity^2^. With the correct choice of *q_ϕ_* and *p_θ_* the right hand side can be evaluated and optimised via gradient descent. Kingma and Welling were the first to recognise that the right hand side has taken on the form of an autoencoder: *q_ϕ_* is *encoding* **x** into **z** and *p_θ_* is *decoding* it back into **x**^3^, see figure 2b.

It is common to assign a unit Gaussian prior distribution to the latent variable, 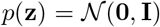. The encoder is also approximated with a multivariate distribution

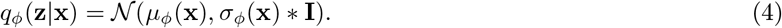

To clarify, the distribution of all latent variables is a unit multivariate Gaussian with zero mean. Once an input is provided, **x**, this collapses to the conditional probability *q_ϕ_*(**z**|**x**), which is a another multivariate Gaussian with means *μ_ϕ_*(**x**) and standard deviations *σ_ϕ_*(**x**). Finally, the conditional probability distribution *p_θ_*(**x**|**z**) is also modelled as a multivariate Gaussian like

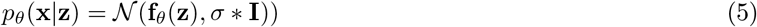

where **f**_*θ*_(**x**) is the decoder neural network and *σ* is a hyperparameter. It is shown in appendix B that the objective function to be *minimised*, i.e. the negative of equation (3), evaluates to:

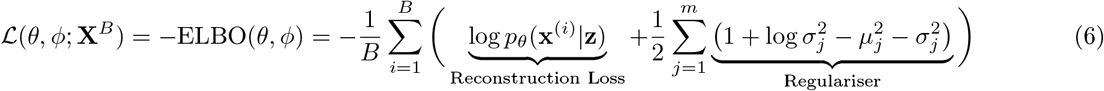

where 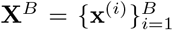 is a minibatch of size *B* drawn randomly from the training data and 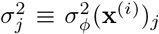. The ‘reconstruction loss’ term encourages the decoder to learn to reconstruct the data well. The second term is known as the ‘regulariser’[34].

#### What benefit do VAEs have over regular AEs?

The issue with regular AEs is that the latent space has little or no structure so the latent variables for very similar inputs can be be completely dissimilar. In VAEs, we forced the latent space to have a continuous Gaussian distribution such that all reasonable samples will produce an output which is similar to values in the training data set. VAEs are therefore extremely good at data generation[35]. Additionally, since we train by sampling randomly from a multivariate Gaussian posterior (equation (4)) the VAE becomes robust against small changes in the input - thus similar inputs are grouped together within the latent space.

In this study a more important consequence it that since the latent space must learn to encode information more carefully, this usually results in the latent variables each encoding highly logical features closely linked to the fundamental physics, biology etc. of the training data^4^. In the context of this report, trained on a data set of ovarian cancer transcriptomes we hypothesise that a VAE might learn to represent platinum resistance in one of its latent variables, entirely unprompted.

### 2.3 t-SNE

t-distributed stochastic neighbour embedding (t-SNE) is a non-linear machine learning clustering algorithm, developed by der Maaten et al. (2008)[37], well suited for embedding high-dimensional data for visualisation in a two dimensional manifold. t-SNE works in a similar fashion to VAEs - first a probability distribution, *p_ij_*, is formed over all pairs high-dimensional data points in such a way that *p_ij_* is large for similar objects and small for dissimilar ones:

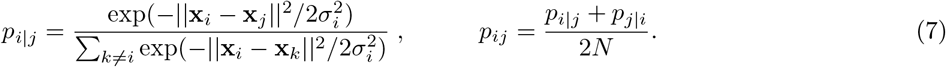

t-SNE then learns a 2-dimensional map {*x_i_*} → {*y_i_*} (with 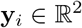) which reflects the similarities in *p_ij_* as well as possible using a Student-t distribution:

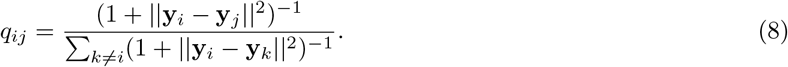

The mapping is optimised, using gradient descent, to minimise the Kullback-Leibler divergence of *q_ij_* from *p_ij_*

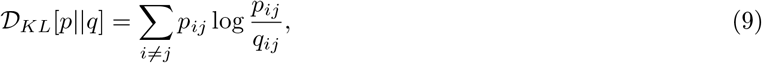

resulting in a collection of points in two dimensions clustered according to the ‘similarity’ of points in the hi0gh dimensional space. In section 4.2 we use t-SNE to further compress our latent space for visualisation in two-dimensions.

## 3 Methods

### 3.1 VAE Architecture

The VAE framework, figure 3, was built in Keras[38], using a Tensorflow[39] backend. Our VAE is modelled on ‘Tybalt’, a VAE designed by Way et al.[21] for a similar although more generalised pan-cancer study. Changes include removing any notion of a ‘warmup’ (shown to be ineffectual), changing the layer activations from *relu* to *sigmoid* and adjusting the input and latent space dimension sizes. The network has one fully connected layer in the encoder (with batch normalisation) and decoder. Training is performed using the AdamOptimiser algorithm[40] to minimise the VAE loss function given in equation (6).

**Figure 3:**
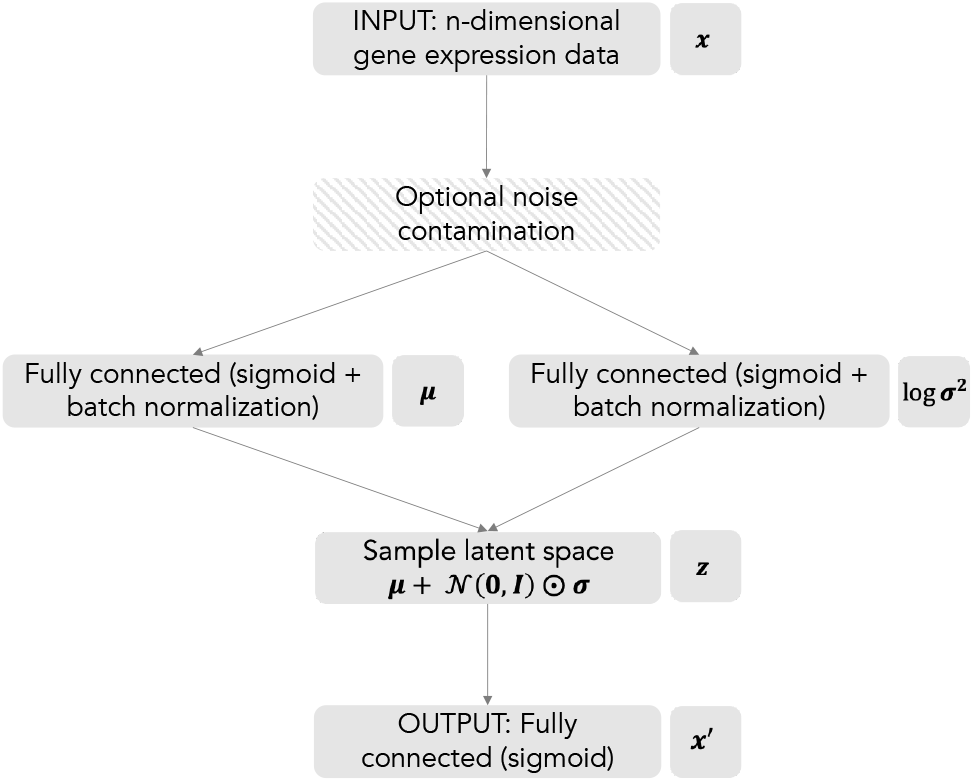
Flowchart illustrating the architecture of our VAE. Each block represents a layer in the network, arrows show the flow of information.

### 3.2 Hyperparameter Analysis

We perform a grid-search parameter ‘sweep’ over all permutations of latent space dimension (5, 10, 20, 50, 100), training epochs (20, 50, 100, 200), batch size (10, 25, 50, 100) and learning rate (0.00025, 0.0005, 0.001, 0.0025) along with 3 variations of the initial data set size (explained in section 3.4), for a total of 960 permutations. The loss was evaluated at each epoch for distinct training and validation sets, these being a random 60:40 partition of the full data set. All training for the hyperparameter sweep was performed in parallel on a remote cluster of 32 AMD Opteron 6376 CPUs at the Computer Laboratory, University of Cambridge. With this set up, a sweep over all 960 hyperparameter permutations took on the order of 30 minutes. The results from 50 such sweeps (~24 hours) were averaged, analysed and plotted in figure 4.

**Figure 4:**
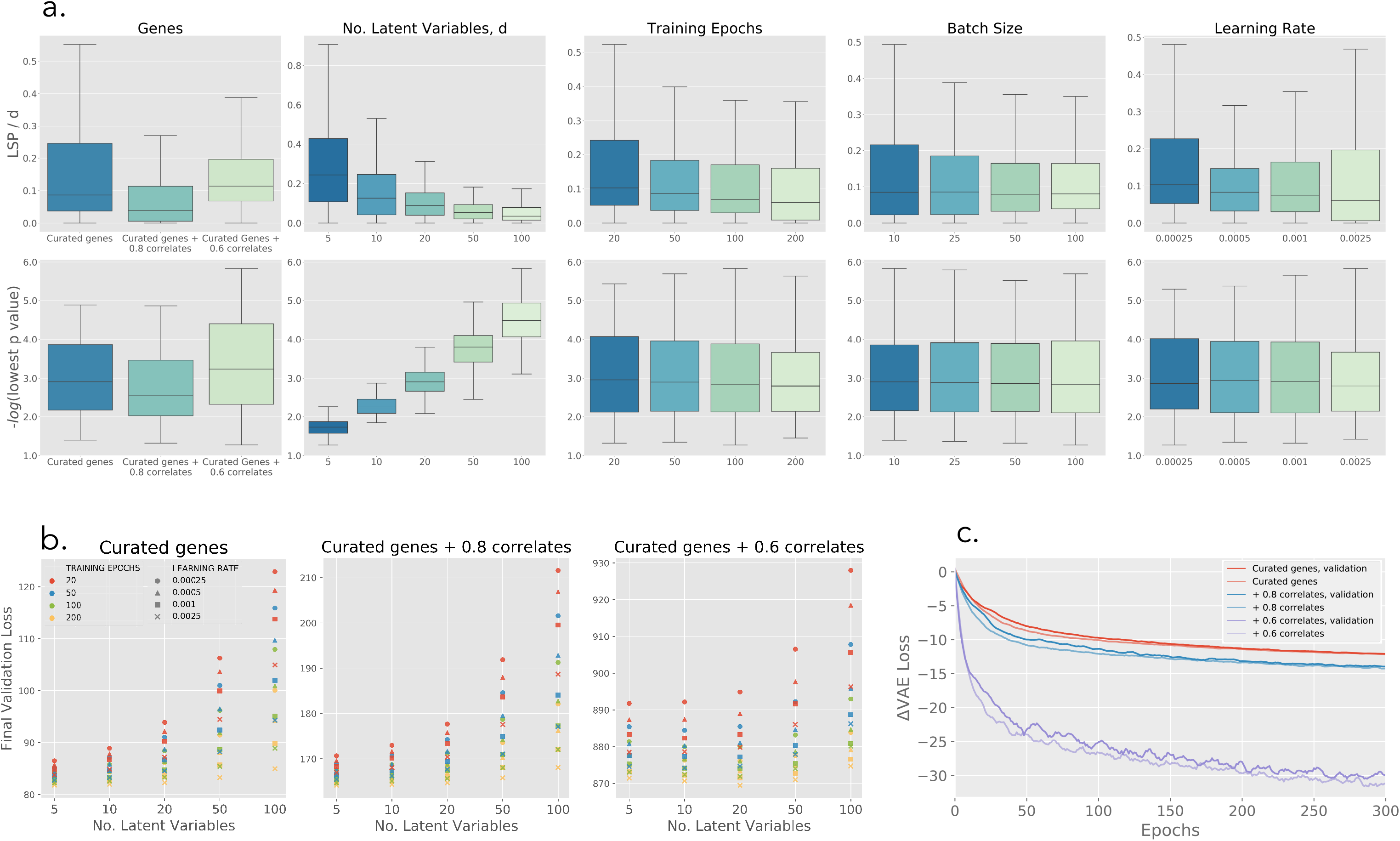
VAE Training and Hyperparameter Analysis: (a) The VAE was tested across a grid search ‘sweep’ of 960 hyperparameter permutations. The plot shows analysis of an average over 50 sweeps. The y-axes gives the test statistic/metric, a high value of either metric is a promising indication that the VAE has learnt a representation of platinum resistance in its latent space, (b) The final VAE loss function is plotted for all hyperparameter permutations (except batch size, which is averaged over), (c) The VAE loss function is plotted over the first 300 epochs (20 latent variables, learning rate of 0.001, batch size of 50). The majority of training is completed within the first 50 to 100 epochs for all three gene sets with relatively little overfitting.

### 3.3 Performance metrics

It is non-trivial to define a single metric which unanimously answers the question: *how good is the VAE at representing platinum resistance in its latent space?* We define a metric hereon referred to as the *Latent Space Performance* or *LSP* as follows:

1. The test data is encoded into the latent space.
2. For each latent variable the resistant and sensitive patients are grouped into two distributions.
3. If the latent variable has learnt to encode platinum resistance we expect these distributions to be significantly different. They are compared using a Kolmogorov-Smirnov test[41] to obtain a p-value for each latent variable.
4. False discovery rate (FDR) is controlled using the Benjamini-Hochberg procedure[42] with the maximum FDR rate set to 25%. This dictates which latent variables are ‘significant’, usually around p < ~ 0.05.
5. The p-values of all ‘significant’ variables are combined into a single metric using Fishers method[43] to give the LSP, equation (10).

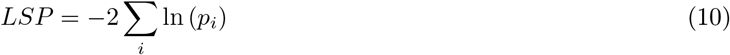

where *i* runs over all latent variables identified as significant in step 4. The LSP can be high, meaning the latent space represents platinum resistance <u>well</u>, for two reasons: either there is one latent variable which *very clearly* distinguishes resistance *or* there are many variables which weakly distinguish resistance, or some combination. Whilst this metric works well as a guide, it is by no means a definitive test for which VAE models are best.

We also use the models lowest p-value as a metric since the corresponding latent variable is highly likely to encode platinum resistance very strongly.

### 3.4 Data

Our data comes from The Cancer Genome Atlas[44] and consists of mRNA expression values for 307 ovarian cancer patients, across (initially) 17,944 genes. Of these, the condition (i.e. resistant or sensitive) is known for just under half. A sample of the data is shown in Table 1.

**Table 1:** A sample of the 0 to 1 scaled mRNA expression data used in this study, obtained from The Cancer Genome Atlas. Each row corresponds to a patient (307 in total), each column corresponds to a gene (17,944 before being reduced for training). The final column corresponds to the patients cisplatin chemotherapy state. The labels are <u>not</u> used for training therefore this is entirely unsupervised machine learning.

|  | ABCE1 | ABCF1 | ABHD13 | ABL1 | ABT1 | ... | Label |
| --- | --- | --- | --- | --- | --- | --- | --- |
| TCGA.36.1581 | 0.5268 | 0.7030 | 0.4195 | 0.4132 | 0.3900 | ... | Sensitive |
| TCGA.09.2056 | 0.3842 | 0.0000 | 0.6427 | 0.2458 | 0.4807 | ... | Sensitive |
| TCGA.25.1318 | 0.4365 | 0.5447 | 0.7401 | 0.5506 | 0.317 | ... | Resistant |
| TCGA.24.2298 | 0.3867 | 0.4281 | 0.8428 | 0.3987 | 0.1915 | ... | Unknown |
| TCGA.31.1953 | 0.3850 | 0.6389 | 0.4361 | 0.3762 | 0.5046 | ... | Resistant |
| TCGA.24.1416 | 0.3117 | 0.2315 | 0.549 | 0.4782 | 0.1372 | ... | Unknown |
| TCGA.25.2398 | 0.2818 | 0.6842 | 0.6444 | 0.6784 | 0.5191 | ... | Sensitive |
| TCGA.25.1027 | 0.2903 | 0.5357 | 0.2388 | 0.8079 | 0.6074 | ... | Resistant |
| ... | ... | ... | ... | ... | ... | ... | ... |

A curated set of 121 genes known to be involved in processes associated with platinum resistance is used to create three variations of the input data defined as follows:

- **Curated genes**: Curated genes only, *x* = [121 genes x 307 patients]
- **Curated genes** + **0.8 correlates**: Curated genes plus all genes which correlate highly (| *ρ* |> 0.8) with any of the curated genes, *x* = [243 genes x 307 patients]
- **Curated genes** + **0.6 correlates**: Curated genes plus all genes which correlate highly (| *ρ* |> 0.6) with any of the curated genes, *x* = [1290 genes x 307 patients]

This is done to reduce the required computational power by starting with a smaller gene set constructed of genes we *expect* to be important^5^. It is a trade off between how likely we are to succeed (addition genes may bring mostly irrelevant information) and how useful we want our models to be after training (if we only use curated genes we can’t expect to infer any <u>new</u> biological pathways). The curated genes are known to be closely related to DNA damage and DNA repair processes in the cell - cisplatin is a DNA damaging agent used in chemotherapy to destroy cancerous cells [45, 46] thus we expect platinum resistance might have something to do with DNA damage/repair.

In section 4.4 we investigate how robust this methodology is to cancer data which has been artificially contaminated with noise. For this task we use a larger TCGA pan-cancer data set of 5000 genes from over 10,000 tumours covering 33 types of cancer. This was done to bypass the issues the comparatively small ovarian cancer data set (which has only 307 tumours) suffers from.

### 3.5 Biological Interpretation of the Latent Space

To infer biological processes which could be linked to platinum resistance we perform gene ontology enrichment analysis for all starting genes which correlate | *ρ* |> 0.2 with any of the significant latent variables. We use g:profiler[47], a web baser server for functional interpretation of gene lists, to perform the enrichment analysis against a large background database of listed gene ontology (GO) terms. Here we focus on biological processes although, in principle, this could be extended to molecular functions, cellular components or other gene sets. The GO term (e.g. G0:0006291) connects a biological process (e.g. DNA Repair) to a list of genes known to be involved in that process - thus by testing for statistical enrichment of these genes in our correlating gene subset we can find biological processes potentially involved in platinum resistance.

## 4 Results and Discussions

The VAE successfully compressed the ovarian cancer data for all three gene sets into a low-dimensional subspace and learnt at least one statistically significant encoding of platinum resistance 17% of the time.

### 4.1 Hyperparameter Analysis

Figure 4a summarises the performance of the VAE across all hyperparameter permutations. Both metrics (the LSP density^6^ and the negative log of the lowest p-value) show that the number of training epochs, the batch size and the learning rate have a relatively weak effect on the performance, with a slight preference toward lower values in all three cases.

The number of latent variables (column 2) has a stronger effect on the performance. Specifically, the LSP density shows a preference towards *smaller* latent spaces - increasingly bottlenecked VAEs are forced to pack more meaningful biological features into fewer latent variables, increasing the LSP density. P-value guided performance shows preference towards *large* latent spaces - models with more latent variables simply have a greater probability of having learn a very strong encoding of platinum resistance in at least one of them. The disagreement between the two plots highlights the underlying challenge in defining an appropriate success metric.

Column 1 shows that performance, by either metric, is slightly better when the selected genes come either only from the curated set *or* from the curated set plus the larger number of correlated genes. Firstly, this confirms that the curated genes *do* have important roles in platinum resistance (they give the best results). Secondly, it shows that adding extra genes outside of the curated set initially makes the results poorer however eventually the VAE responds to the presence of additional genetic information by more selectively encoding platinum resistance.

The final validation loss is affected strongly by the choice of hyperparameter (Figure 4b). Generally as the number of latent variables is increased the final validation loss increases and deviates more significantly for each hyperparameter. The majority of the VAE training occurs in the first 50 epochs (Figure 4c) although after 300 epochs the loss function is still slowly decreasing and relatively little overfitting is observed. We argue it is not necessarily problematic that some of the hyperparameter permutations have their training terminated before saturation since our primary goal is not to find models which minimise some loss function, but instead models which distinguish platinum resistance. Thus it is OK to treat the number of training epochs as a hyperparameter, similar to Way et al.[21]. We elaborate on this in section 4.4.

On one of the 50 hyperparameter sweeps *all* 960 models were saved for downstream analysis. Table 2 identifies nine high performance models which we use for analysis in later sections.

**Table 2:** Nine VAE models identified as high performers and are listed here along with their hyperparameters and metric values. *A* and *B* models both are discussed in section 4.2. *C* models are used to generate genes for the enrichment analysis as discussed in section 4.3. Bold values illustrate the metric used to choose the model.

| Model Code | Genes | Latent Space Dimension, d | Training Epochs | Batch Size | Learning Rate | LSP/d | Best p-value | Significant Variables |
| --- | --- | --- | --- | --- | --- | --- | --- | --- |
| A1 | Curated | 50 | 100 | 25 | 0.00050 | 0.29 | **0.00077** | 1 |
| A2 | +0.8 Correlates | 50 | 50 | 50 | 0.00025 | 0.45 | **0.00001** | 1 |
| A3 | +0.6 Correlates | 100 | 20 | 25 | 0.00100 | 0.19 | **0.00009** | 1 |
| B1 | Curated | 5 | 50 | 10 | 0.00250 | **2.16** | 0.05219 | 2 |
| B2 | +0.8 Correlates | 5 | 50 | 50 | 0.00025 | **2.23** | 0.00384 | 1 |
| B3 | +0.6 Correlates | 10 | 100 | 50 | 0.00250 | **2.10** | 0.01974 | 3 |
| C1 | +0.6 Correlates | 5 | 100 | 10 | 0.00250 | **2.17** | 0.05081 | 2 |
| C2 | +0.6 Correlates | 10 | 200 | 25 | 0.00250 | **2.34** | 0.01643 | 3 |
| C3 | +0.6 Correlates | 100 | 100 | 100 | 0.00025 | 0.14 | **0.00109** | 1 |

### 4.2 Latent Space Interpretation

Models A1, A2 and A3 from table 2 all had one latent variable which learned an encoding representing platinum resistance. The p-values for these significant variables are 2 to 4 orders of magnitude smaller than the next best latent variable (note the log scale in the lower panel of figure 5), thus we can be fairly sure this is not a ‘fluke’ but is indeed a biologically meaningful feature the VAE has learned during training. It is possible that platinum resistance can be encoded into more than one latent variable. In model A3, for example, VAE dimensions 12 *and* 90 had somewhat lower p-values than the rest of the dimensions but only dimension 12 was classified as significant. Only a small proportion (~17%) of VAEs learnt a statistically significant encoding of platinum resistance, of which only a handful were as clear as the models visualised in figure 5. This likely implies that the signal or ‘imprint’ of platinum resistance is relatively weakly encoded into the transcriptomes.

**Figure 5:**
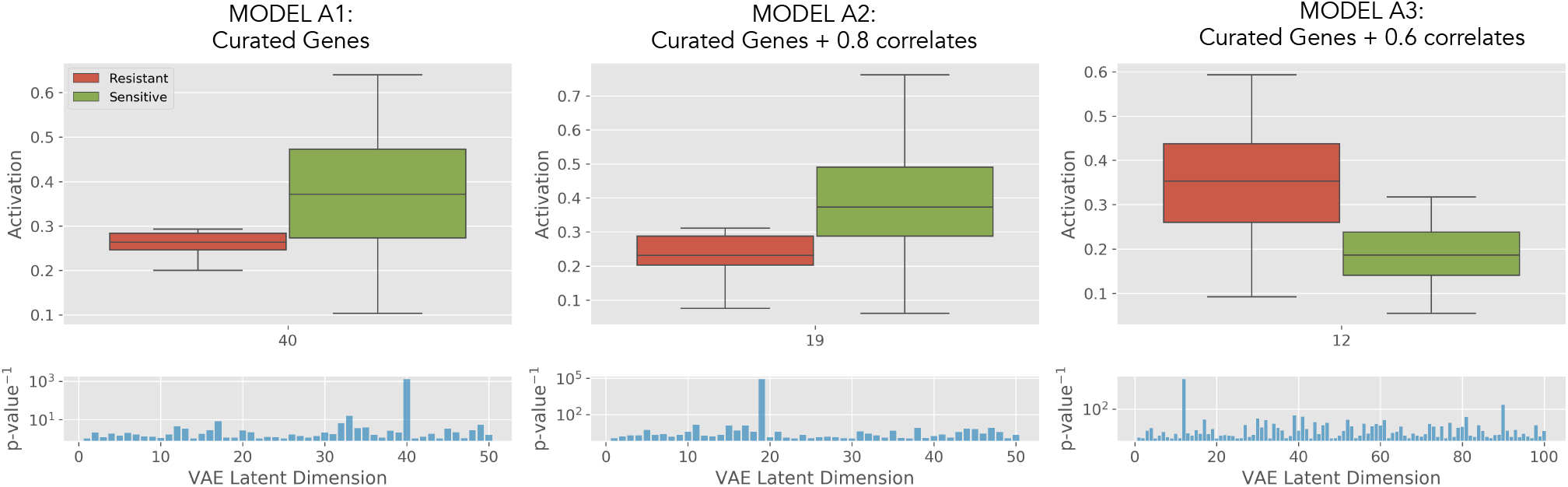
Variation autoencoders trained on TCGA ovarian cancer data learn latent features representing platinum resistance. The lower panel displays the Kolmogorov-Smirnov p-value of *all* latent variables. The Kolmogorov-Smirnov test asks whether the distribution of resistant patients is significantly different from the distribution of sensitive patients (read as: asks whether the latent variable has learnt to represent platinum resistance).

One issue which over arches this study is the relative smallness of the data set we are using. Compared to Way’s study[21] which uses a pan-cancer data set of over 10,000 tumours our data set has only 307. Furthermore, not all data comes with resistance labels and we use 60% of the data for training, meaning that there are only 15 resistant patients and 45 sensitive data points with which to produce figures 5 and 6. Unfortunately there is currently little that can be done: the 45:15 split is a biological feature of how the cancer manifests itself and the TCGA data set we used is the largest publicly available ovarian cancer transcriptome data set with matched clinical data. With that said, the signals we see here *are* promising and suggest similar results could be validated in an adequately powered data set, were one to exist.

**Figure 6:**
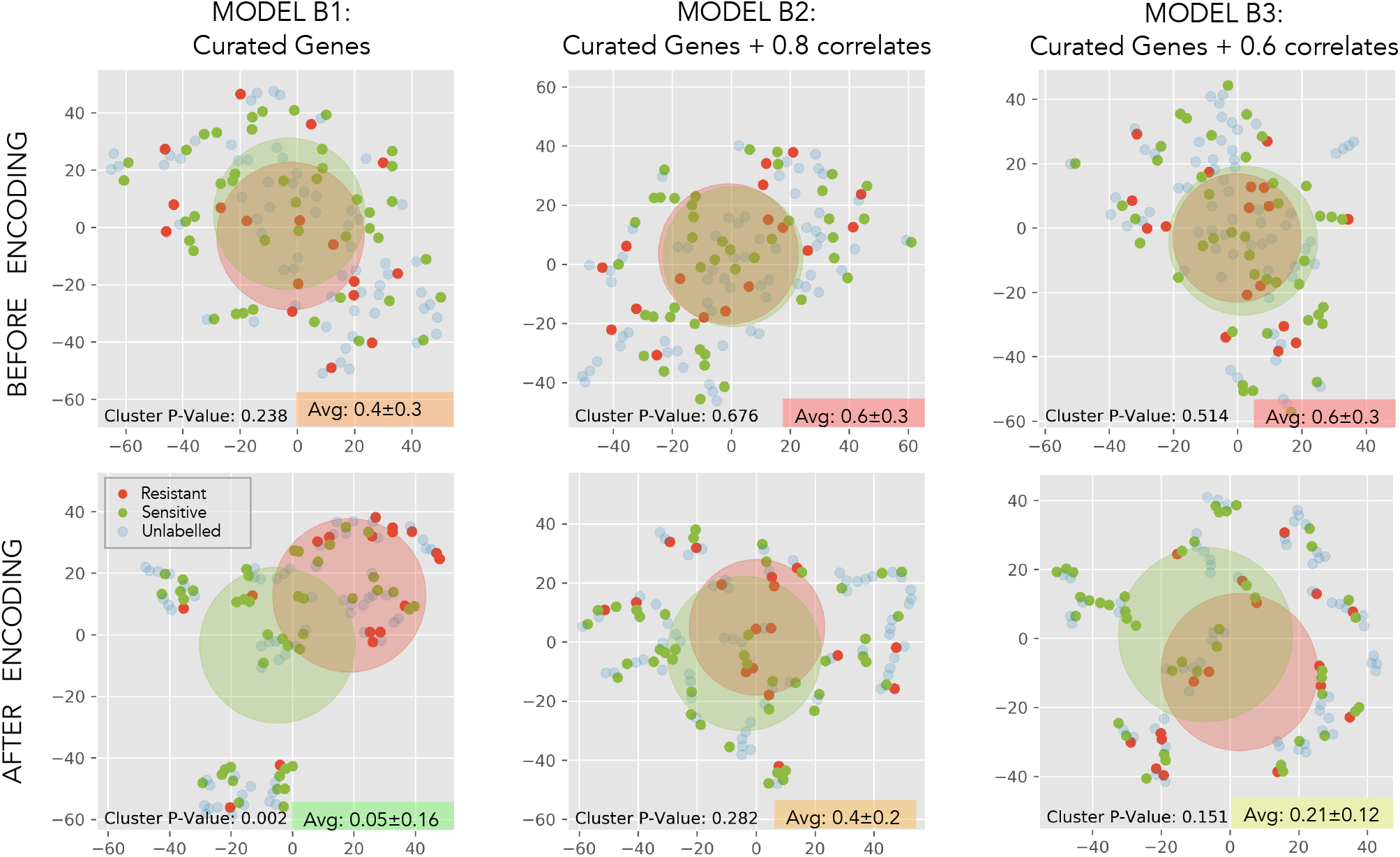
t-SNE clusterings of ovarian cancer gene expression data before and after VAE encoding. P-values are for a two-dimensional two-sample Kolmogorov-Smirnov test on the distributions of resistant and sensitive patients. Averages p-values over 30 clustering attempts are shown.

t-SNE was used to cluster the data before and after VAE encoding into a two-dimensional manifold for visualisation, for models B1, B2 and B3. These models had a high LSP density and so we hypothesised that t-SNE might be able to cluster resistant/sensitive patients together - at least to a better extent than for the raw data. Figure 6 partly confirms this hypothesis. By visualising the mean and standard deviation of the two patient groups with faded circles, one can see that they have been separated by t-SNE *only* after encoding. This is confirmed by the Kolmogorov-Smirnov test p-values which shows, in all three case, that the 2D distribution of resistant and sensitive patients are more distinct (the p-value has decreased). *However* it must be pointed out that in only one of the models shown (B1) was the distribution separation significant (*p* ≤ 0.05) when averaged over 30 t-SNE clustering attempts.

t-SNE has a very hard job clustering the raw data (the algorithm returns a “ball” of approximately uniformly distributed points) but *after* encoding clusters begin to appear. This demonstrates that the information in the latent space is less noisy than the input data and has probably been distilled into more meaningful and robust biological features which t-SNE can use to group patients with similar genetic traits. This figure gives us visual proof that VAEs significantly improves the ability of t-SNE to cluster data points and that, in a handful of models, the latent space is sufficiently concentrated with platinum resistance information that the two patient groups are distinguishable (if not quite ‘clustered’) using t-SNE.

### 4.3 Enrichment Analysis

Correlations between significant latent variables and the original gene set are used to extract subsets of genes which are potentially related to the biological processes we cumulatively call platinum resistance. This was done for three models; C1, C2 and C3. The three genes lists contained 96, 85 and 67 genes respectively and the overlaps, shown in figure 7a, are significantly larger than one would expect for three randomly selected subsets. This strongly suggests the three subsets are biologically linked, as we are hoping they would be. It is promising to see BRCA2, a gene well known to be linked to ovarian cancer[48], independently appearing in two of our three gene subsets.

**Figure 7:**
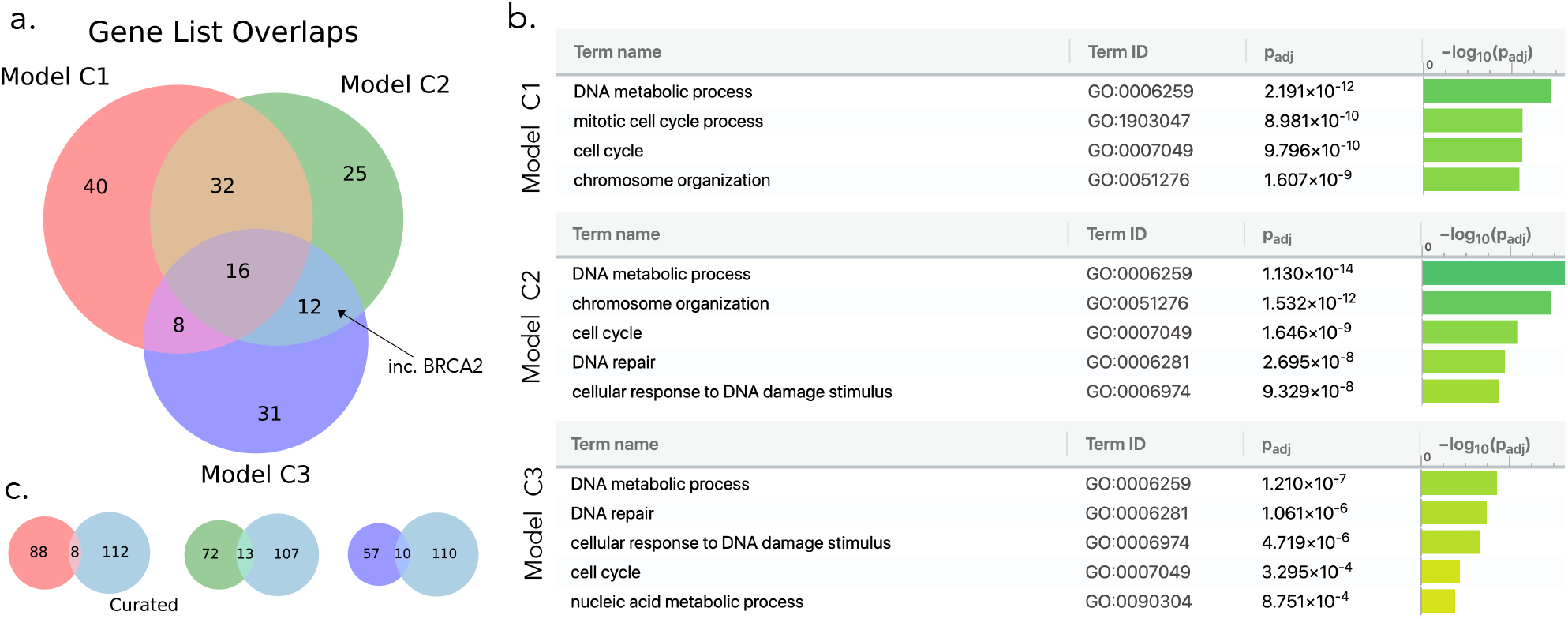
A summary of gene ontology enrichment analysis performed on genes subsets extracted from platinum resistance distinctive latent spaces. Genes were selected if they correlated | *ρ* |> 0.2 with any of the significant latent variables. (a) Overlaps for gene subsets collected from three distinct models. (b) A summary of the most enriched gene ontology terms for each model. All models point towards similar biological processes being involved in platinum resistance - these largely revolve around DNA repair and cell response to DNA damage. (c) Overlaps of the three gene subsets with the curated gene list.

Figure 7b shows the most significant gene ontology terms for each of the three gene subsets. All models shared the cell cycle GO term (GO:0007049), whilst models C2 and C3 identified DNA repair (GO:0006281) and DNA damage (GO:0006974) GO terms. One possible reason for this is that the initial ‘curated’ gene list itself contains genes closely linked to DNA damage (as explained in section 3.4). However, as shown in figure 7c, the overlap of the three extracted gene sets with the curated gene list is are not overly large, suggesting what we are seeing here might be independent verification that DNA damage and DNA repair processes are closely linked to platinum resistance.

### 4.4 Response of VAE training to noisy cancer data

The VAE was trained to compress the larger pan-cancer data set (of 5000 genes across 10000 tumours) into a latent space of 100 dimensions. Gaussian and dropout noise was added separately to the data before encoding. The decoder was then tasked to reproduce the *clean* input from the latent encoding of the contaminated input. The noise fraction is defined as the standard deviation of the added Gaussian noise divided by that of the original data, or in the case of dropout noise, the proportion of the genes deleted divided by the proportion remaining. Figure 8 shows the VAE loss (equation (6)) and the reconstruction error (equation (1), averaged over all patients and genes) after 50 epochs of training. In both plots a *lower* value implies *better* training.

**Figure 8:**
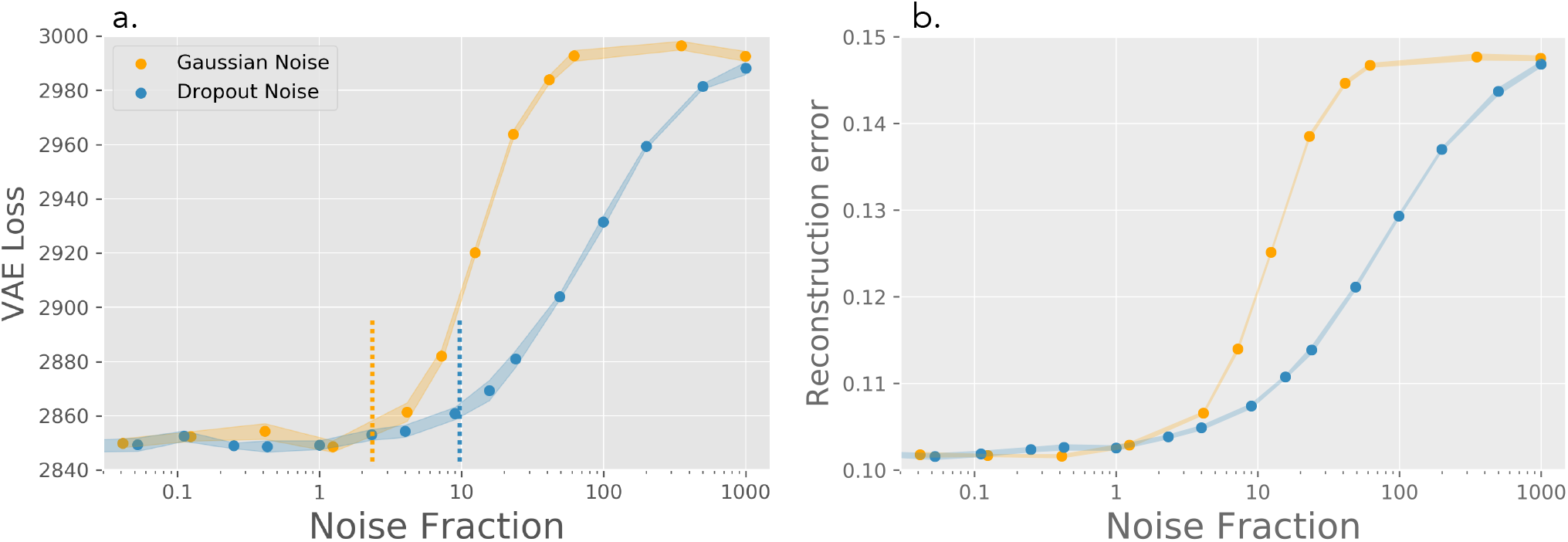
VAE response to noisy cancer data. (a) The VAE loss function evaluated after 50 epochs. (b) The average euclidean reconstruction error per gene evaluated after 50 epochs. Noise fraction refers to the proportion of the input data which can be accounted for by artificially added noise. Shaded regions display approximate error bounds (standard deviation over 6 runs). Dashed lines denote approximate noise fraction where result begin to deteriorate.

The VAE training is almost entirely unaffected by the addition of Gaussian noise up to a noise fraction of between 2 and dropout noise up to a fraction of 10. This confirms the common knowledge that autoencoders, which must squeeze only the most *essential* information through the bottleneck, are in general, excellent at denoising a signal. It is pleasing to see this applies to cancer gene expression data, which due to the inherent complexity of making genome microarray measurements, is often noisy and/or lossy.

Beyond a noise fraction of 100, both loss functions plateau. This is where the input is entirely dominated by noise and so the VAE suffers complete failure - essentially it can do no better than predict a constant mean value for each gene in the decoding. This corresponds to a reconstruction error of 0.15 (figure 8b), only 50% higher than the reconstruction error for clean data posing the question “was the reconstruction good to begin with?”. The answer is, to the human eye, no: an average reconstruction error of 0.1 per gene (for data initially normalised between 0 and 1) is fairly large. However, unlike for a regular autoencoder, reconstruction of the input is *not* the primary function of a VAE and, as demonstrated by Way et al., meaningful biological features can be encoded in the latent space even if the decoder is then relatively bad at reconstructing the gene expression values afterwards. The same should hold for the ovarian cancer data used in other parts of this study.

## 5 Conclusions

We trained a variational autoencoder on a data set of ovarian cancer transcriptomes and showed that, in many cases, one or more of the latent variables learnt an encoding which represents platinum resistance. From here it was possible to perform enrichment analysis on genes correlating highly with the distinguishing latent variables and identify biological processes closely linked to platinum chemotherapy resistance. We showed that t-SNE could be an effective tool at visualising the encoding of platinum resistance within the latent space however, in the majority of models, the signal was not strong enough statistically confirm this result. Finally, we showed that the VAE is highly robust to cancer data contaminated with large amounts of Gaussian and dropout noise, an important feature if this method is to be properly validated on larger ovarian cancer data sets *or* applied to problems in non-medical fields.

In the majority of attempts no clearly significant encoding of platinum resistance was found in the latent space identifying the two main limitation of this study: firstly, it appears that at a genetic level platinum resistance does not impart a strong enough signal on the transcriptome for the VAE to learn an encoding every time. Secondly, the data set is small and so our results varied a lot run-to-run. With a larger (say, 10x larger) data set we would expect more consistent and reproducible results from which we could draw firmer conclusions. Nonetheless, we believe these result show promising, if not quite conclusive, signs that VAEs could be a robust way to investigate poorly understood features in cancer from an entirely unsupervised and data science oriented viewpoint. However, due to the practical issues of data set size, it is currently only fair to say that this requires more careful validation and evaluation.

Crucially this is an exercise in data science applied to the field of cancer research, <u>not</u> the other way round. The entirely unsupervised nature of this research means it could almost immediately be adapted to other fields, for example physics, where there is a desire to extract novel insight from complex, high-dimensional and potentially very noisy data.

## 6 Reproducibility

All data and code (in the form of an easy to use Jupyter notebook) has been made available on Github at the following address: https://github.com/TomGeorge1234/oVAErian-Cancer.

## 7 Acknowledgments

The authors gratefully acknowledge support from Dr Syed Haider (London Institute for Cancer Research) for his significant and continual contributions to this work.

## A Deriving Variational Bayes

The Kullback-Leibler divergence is defined as

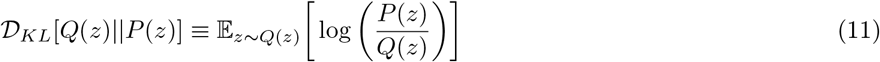

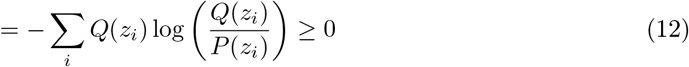

where equality is achieved iff *Q* ≡ *P*. Let’s evaluate the KL divergence between the distribution we will sample **z** from, *q_ϕ_*(**z**|**x**), and the true posterior *p*(**z**|**x**)

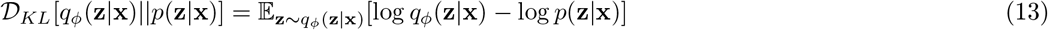

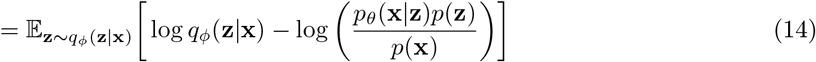

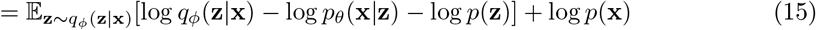

where in the second line we have used Bayes’ law and in the third we pull log *p*(**x**) out from the average since it is indepent of **z**. Rearranging give the required result, equation (3)

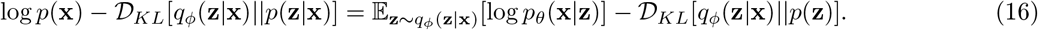

## B Evaluating the VAE objective function

Evaluating the second term on the right-hand side of equation (3) is easy since the Kullback-Leibler divergence between two multivariate Gaussians is a known quantity, see [49]

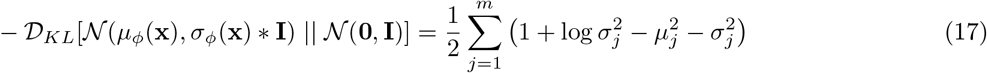

where 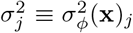, and the same for *μ*. The first term is not so easy to evaluate. It can be approximated using Monte Carlo as

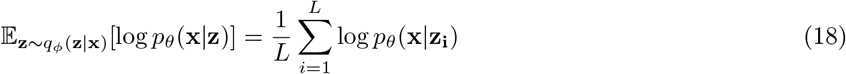

where **z** is sampled from *q_ϕ_*(**z**|**x**). It turns out (see Kingma and Welling [29]) that if the batch size is large enough, then *L* = 1 is a big enough sample size. This means on each forward pass of the variational autoencoder we only need to sample from the posterior of a given **x** in the batch once, significantly speeding up the training process. This Monte Carlo sampling is represented by the four central arrows in figure 2b. Putting these terms together gives the full objective function, as given in equation (6).

## Footnotes

1 The theory and code provided ought to be sufficient that this methodology could quickly be adapted and applied to a more classically “physics” oriented research problem.

2 i.e. if our model for *q_ϕ_*(**z**|**x**) is sufficiently general, it will rapidly approximate the true posterior *p*(**z**|**x**) during training and this error should become negligible.

3 Although figure 2b shows a latent vector being mapped directly to the output, **x**′, we still say the decoder is learning the probability distribution for **x**′, not **x**′ itself. What the decoder network is actually outputting is the multivariate Gaussian means as per equation (5). This distinction can be confusing, whereas an autoencoder maps **x** → **z** and then **z** → **x** directly, the variational autoencoder maps **x** → *q_ϕ_*(**z**|**x**) which we can sample, then pass through the decoder to map **z** → Mean [*p_θ_*(**x**|**z**)].

4 For example, training on a broad data set of human face images, the latent variables of a sufficiently adept VAE will learn to smoothly control a persons skin colour, facial hair, or, whether or not they’re wearing glasses[36].

5 Note: we are <u>not</u> claiming we already know which genes/gene pathways cause platinum resistance - if that was the case this study would be redundant. We are merely making a ‘best guess’ based on our current understanding, to slim down the initial 17,944 genes (most of which will be entirely unimportant in platinum resistance) to a more informed subset.

6 i.e. LSP per latent variable, the motivation being that if the LSP density is high the latent space is “dense” in platinum resistance information which is desirable

